# Nucleus remodeling activity is conserved amongst diverse SIV Vpr isolates

**DOI:** 10.64898/2026.09.02.749000

**Authors:** Zaddock Williford, Emily Leavitt, Nicholas Saladino, Madisyn Johnson, Daniel J Salamango

## Abstract

Human immunodeficiency virus (HIV) encodes four accessory proteins that are essential for virus replication *in vivo*, primarily through the counteraction of host innate immune defense mechanisms. One of these proteins, Vpr, induces constitutive DNA damage repair (DDR) signaling to drive global epigenetic remodeling and activation of transcription programs that enhance HIV-1 promoter activity during acute infection and virus reactivation from latency. Vpr is conserved amongst diverse simian immunodeficiency virus (SIV) strains; however, the evolutionary breadth of Vpr’s nucleus remodeling activity has yet to be thoroughly characterized. Here, we investigate a diverse panel of 16 SIV Vpr isolates and demonstrate that 13 out of 16 are capable of significantly activating DDR signaling compared to control infected cells. Moreover, cells infected with these isolates also exhibit increased abundance of two histone marks associated with transcription and euchromatin formation, as well as increased activation of two transcription factors known to be critical for HIV-1 promoter activity. Furthermore, site-directed mutagenesis of a highly homologous SIV Vpr isolate that failed to engage the DDR response revealed previously uncharacterized amino acid residues required for HIV-1 Vpr DDR engagement. Finally, structural modeling and functional analyses revealed that phylogenetically diverse Vpr isolates from SIV African green monkey strains induce nucleus remodeling through an evolutionarily distinct set of amino acid residues. Together, these findings demonstrate that hijacking of DDR responses to promote remodeling of the nuclear environment is a broadly conserved Vpr function.

## INTRODUCTION

Human immunodeficiency virus type 1 (HIV-1) derives its evolutionary origins from a closely related group of lentiviruses that infect non-human primates (simian immunodeficiency viruses [SIVs]). Five distinct lineages of primate lentiviruses have been identified and are attributed to at least seven cross-species transmission events that have given rise to HIV-1 Groups M, N, O, and P, as well as HIV-2 (Sharp and Hahn, 2010). HIV and SIV lineages encode similar gene sets, including several accessory proteins that are dispensable for virus replication *in vitro* but are essential for viral pathogenesis *in vivo*. One such accessory protein, Vpr, is highly conserved amongst HIV-1 and several SIV strains (including chimpanzee, CPZ; gorilla, GOR; and red-capped mangabey, RCM) but is more divergent amongst HIV-2 and ancestral SIV strains (such as sooty mangabey, SMM; and African green monkey, AGM) (**Figure 1A**).

**Figure 1.**
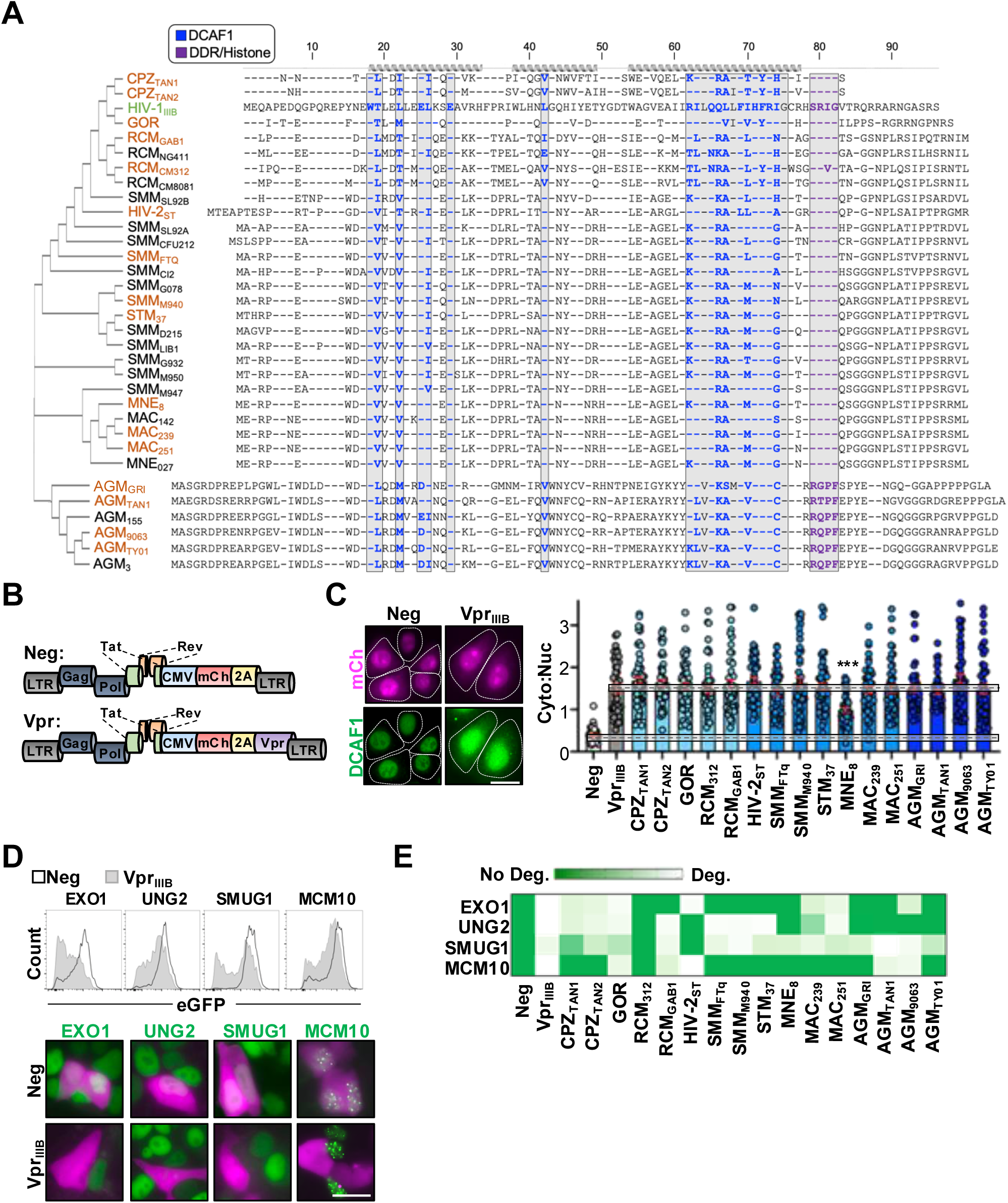
Functional validation of SIV Vpr isolates. (**A**) Amino acid alignment of the indicated Vpr isolates. SIV Vpr sequences were obtained from the Los Alamos database and are stratified phylogenetically based on the HIV-1_IIIB_ Vpr sequence (green). Isolates colored in orange were investigated in this study, with amino acid residues required for DCAF engagement colored in blue and amino acid residues required for induction of DDR signaling, epigenetic remodeling, and transcription activation are colored in purple. Dashed lines represent amino acid residues that are conserved relative to HIV-1_IIIB_ Vpr. CPZ, chimpanzee; GOR, gorilla; RCM, red-capped mangabey; SMM, sooty mangabey; STM, stump-tailed macaque; MNE, pig-tailed macaque; MAC, rhesus macaque; AGM, African green monkey. (**B**) Schematic of proviruses used in this study. The Nef open reading frame has been replaced with a CMV-*mCherry-T2A* cassette which allows for equal expression of all Vpr isolates in addition to HIV-1 genes necessary for infection establishment. (**C**) Representative immunofluorescence microscopy images (left, scale bar = 20 μM) and quantification (right) of DCAF1 localization in HeLa cells infected with the indicated viruses. Cells were infected for 48 hours with the indicated virus prior to being subjected to immunofluorescence microscopy evaluating endogenous DCAF1 localization. Dashed outlines overlaid on the representative images highlight cell boundaries to emphasize DCAF1 relocalization to the cytoplasm. Dashed boxes overlaid on the quantification highlight the mean and standard error mean of DCAF relocalization in control or HIV-1_IIIB_ Vpr infected cells (n = 50). Analyses performed using a one-way ANOVA; ***, p < 0.001. (**D**) Flow cytometric analysis (top) and representative fluorescence microscopy images (bottom) of HEK293T cells stably expressing the indicated eGFP-tagged substrate. Cells were infected for 48 hours with control or HIV-1_IIIB_ Vpr expressing virus prior to being subjected to fluorescence microscopy and flow cytometry. Scale bar = 10 μM. (E) Substrate degradation activity of SIV Vpr isolates against the indicated HIV-1_IIIB_ Vpr substrates. The eGFP-tagged cell lines depicted in (**D**) were infected for 48 hours with the indicated viruses prior to being subjected to flow cytometry to quantify the amount of substrate depletion (*i.e.,* quantifying mean eGFP intensity in mCherry positive cells). Mean eGFP fluorescence was normalized to control and HIV-1_IIIB_ Vpr infected cells to generate the color-coded heat map.

It has been well established that Vpr is essential for virus replication *in vivo* and in macrophages, but the molecular mechanism(s) underlying Vpr’s proviral function remain enigmatic (Connor et al., 1995; Eckstein et al., 2001; Lang et al., 1993; Nodder and Gummuluru, 2019). Vpr’s best characterized activity is the ability to induce systems-level remodeling of the host transcriptome and proteome by hijacking a host E3-ubiquitin ligase complex to target over three dozen cellular proteins for proteasomal degradation (Le Rouzic et al., 2007; Wen et al., 2007). Recent quantitative proteomics studies have revealed that many of these substrates are involved in DNA repair or DNA modification, which rationalizes longstanding observations that Vpr induces constitutive DNA damage repair (DDR) signaling and G2/M cell cycle arrest (Fabryova and Strebel, 2019; Greenwood et al., 2019). In addition, Vpr enhances HIV promoter activity, reactivates proviruses from latency, and directly engages DNA to alter conformational states and potentially induce DNA damage (Fabryova and Strebel, 2019; Iijima et al., 2018; Johnson et al., 2022; Lai et al., 2005; Levy et al., 1994; Levy et al., 1995; Li et al., 2020; Lyonnais et al., 2013; Romani et al., 2016; Zhang and Bieniasz, 2020; Zhang et al., 1998; Zhou et al., 2021). Moreover, in limited testing Vpr-induced DDR signaling has been observed in some HIV-2 and SIV isolates, further suggesting an importance for virus replication (Li et al., 2020). These observations are in line with a growing body of evidence that demonstrate the hijacking of DDR responses to facilitate virus replication is a broadly conserved strategy used by diverse viral families (Saladino and Salamango, 2024).

Recent work from our group has established a direct cause-and-effect mechanism wherein Vpr-induced DDR signaling promotes epigenetic remodeling and activation of transcriptional programs that enhance HIV-1 promoter activity during acute infection and virus reactivation from latency (Saladino N, 2026). Here, we expand on these initial observations by testing an extensive panel of Vpr isolates from diverse SIV species to assess the evolutionary breadth of nucleus remodeling activity. We demonstrate that SIV Vpr isolates phylogenetically similar to HIV-1 exhibit robust DDR modulation activity, while isolates phylogenetically similar to HIV-2 vary greatly in their capacity to induce DDR responses. Diverse SIV Vpr isolates induce activation of DNA repair kinases ATM and ATR, increase the abundance of two histone marks associated with transcription, and activate HIV-associated transcription factors at comparable levels to that of HIV-1 Vpr. Additionally, using structure-guided mutagenesis of an SIV Vpr isolate from red capped mangabey we identify a previously uncharacterized set of amino acid residues used by HIV-1 Vpr to induce DDR signaling. Finally, we investigate the ability of Vpr isolates from

African green monkey SIV to induce robust DDR responses even though they share virtually no amino acid similarity to that of HIV-1 Vpr. Unsurprisingly, structure-guided mutagenesis of these AGM isolates revealed a unique set of Vpr amino acid residues required for inducing DDR signaling, highlighting the remarkable evolutionary capacity of viral accessory proteins to adapt to their host environment.

## MATERIALS AND METHODS

### Cell culture conditions, SIV Vpr sequence analysis, and provirus construction

HEK293T and HeLa cells (American Type Culture Collection) were maintained in DMEM medium (Gibco cat #11-965-118) supplemented with 10% fetal bovine serum (FBS; Gibco, Gaithersburg, MD) and 0.5% penicillin-streptomycin (50 units; Gibco, Gaithersburg, MD). Vero E6 cells (American Type Culture Collection) were maintained in MEM medium (Gibco cat #61-100-061) supplemented with 10% fetal bovine serum (FBS; Gibco, Gaithersburg, MD) and 0.5% penicillin-streptomycin (50 units; Gibco, Gaithersburg, MD). THP1 cells (American Type Culture Collection) were maintained in RPMI medium (Gibco; cat #11-875-093) supplemented with 10% FBS and 0.5% penicillin-streptomycin. For THP1 differentiation, cells were incubated with 100 ng/ml of phorbol 12-myristate 13-acetate (PMA; Sigma #P8139) for 48 h. For generating virus stocks, HEK293T cells were co-transfected with a VSV-G expression vector along with the indicated proviral plasmid. Medium was collected 48 h post-transfection and frozen at minus 80 degrees C until used. For generating eGFP-tagged stable cell lines, substrates of interest were cloned into a pQCXIH retroviral expression construct and co-transfected along with a MLV GagPol packaging plasmid and VSVg expression plasmid in HEK293T cells. 24 hours post-transfection culture media was exchanged with fresh DMEM and viral supernatants were harvested 48-hours post transfection. Samples were purified through 0.22 μM syringe filtration and overlaid onto fresh HEK293T cells. After 48-hours, pure cell populations were generated by treating with 200 μg/mL hygromycin B (Fisher, AAJ6068103).

Available SIV Vpr amino acid sequences were downloaded from the Los Alamos sequence repository (https://www.hiv.lanl.gov) and compared to the HIV-1_IIIB_ Vpr reference sequence as depicted in **Figure 1A**. Amino acid alignments were generated using the publicly available Clustal Omega software.

Proviral expression plasmids have been described previously (Saladino N, 2026). Briefly, all SIV *Vpr* genes were ordered a gblocks from Integrated DNA Technologies (IDT) and cloned into the *mCherry-T2A* expression cassette using *XmaI* and *XhoI* restriction enzymes. The same cloning strategy was used to insert HIV-1_IIIB_ Vpr while the empty vector serves as the mCherry control.

### Fluorescence microscopy and immunostaining

For immunofluorescence microscopy experiments using HeLa cells, approximately 5,000 cells were seeded into a 96 well glass-bottom imaging plate (Ibidi #89627) and allowed to adhere overnight. The next day, cells were infected with the indicated virus for 48 h prior to being washed 1x with PBS and then fixed using 4% paraformaldehyde (PFA) for 10 min at room temperature. After fixation, cells were washed 3x with PBS in 5-minute intervals and permeabilized using PBS plus 0.3% Triton X-100 (PBST) for 10 minutes at room temperature. Cells were blocked using PBST supplemented with 5% bovine serum albumin (BSA, Fisher Bioreagents BP9703100), 10% goat serum (Sigma-Aldrich, G9023-10mL) and 0.3 M glycine for 2 hours at room temperature while rocking. After blocking, samples were incubated with primary antibody against DCAF1/VprBP (1:400, Proteintech 11612-1-AP) in blocking buffer overnight at 4 degrees C. The next day, cells were washed 3x with PBS in 5-minute intervals and then incubated with anti-mCherry conjugated to Alexa Fluor 594 (1:800, Invitrogen M11240) and secondary anti-rabbit-IgG conjugated to Alexa Fluor 488 (1:800, Cell Signaling 4412) in blocking buffer at room temperature for 1 h. After incubation, cells were washed 3x with PBS at 5-minute intervals and stained with NucBlue stain (Thermo Fisher, R37605) and imaged. An EVOS M500 fluorescence microscope was used for imaging, using 60x or 100x oil-immersion objectives.

For immunofluorescence microscopy experiments using differentiated THP1 cells, approximately 50,000 cells were seeded into a 96-well glass-bottom imaging plate in the presence of 100 ng/ml PMA and co-infected with the indicated virus for 48 h prior to being subjected to immunofluorescence microscopy using primary antibodies against γH2A.X (1:300, Cell Signaling 9718), acetyl-histone H3K9 (1:1000, Cell Signaling 9649), acetyl-histone H3K14 (1:1000, Cell Signaling 7627), NF-κB p65 (1:300, Cell Signaling 82425S), c-Jun (1:400, Cell Signaling 9165T), or Phospho-c-Jun (Ser73) (1:800, Cell Signaling 3270T).

### Inhibitor treatments

For inhibitor experiments, cells were treated 24-hours post-infection with either 10 nM ATM inhibitor (Fisher, #AZD1390), 10 µM ATR inhibitor (Fisher, #NU6027), or combination ATM/ATR inhibitor treatment for 24 hours. Cells were then processed for downstream analysis as indicated.

### Flow cytometry

At 48 h post-infection, cells were washed 1x in cold PBS, harvested following trypsinization, and centrifuged at 500 RCF for 10 minutes. After washing 1x with cold PBS, cells were centrifuged and resuspended in 4% PFA for 30 minutes, washed 3x with PBS, resuspended in cold PBS, and subjected to flow cytometry using a Becton Dickinson LSR II flow cytometer.

### Experimental replicates, statistical analyses, and protein modeling

All experimental procedures, except flow cytometry experiments, were repeated at least two times by two independent investigators (n = 4). All flow cytometry experiments were repeated 3 independent times, and the data were analyzed using FlowJo v10 software. For immunofluorescence microscopy experiments, representative data are depicted from one of four experimental replicates (n = 50 cells per experiment). Focus formation and mean fluorescence intensity (MFI) analyses were calculated using ImageJ software and analyzed using GraphPad Prism 6 software. Briefly, boundaries of infected cell nuclei were defined using DAPI staining as an indicator and then foci were quantified using the “find maxima” feature and eGFP mean fluorescence intensity was defined by analyzing integrated pixel intensity of the defined nuclear area minus the background signal intensity of an adjacent area with identical dimensions. Foci prominence thresholding was set so that uninfected negative controls yielded minimal foci. Statistical analyses were performed using GraphPad Prism 8 software after confirming that all data followed a normal distribution. AlphaFold 3 was used to generate predicted structures of the human DCAF1 helix-loop-helix motif and WD40 domain (Ala1044-Glu1398) and HIV-1 subtype B, AGMGRI, or RCMGAB1 Vpr. The top-ranked structure for each complex was aligned to DCAF1 in the DDB1-DCAF1-Vpr-UNG2 co-structure (PDB: 5JK7). The ipTM/pTM confidence intervals for these putative Vpr-DCAF co-complexes were 0.85/0.85, 0.87/0.88, and 0.67/0.7 for HIV-1, RCM_GAB1_, and AGM_GRI_ Vpr-DCAF co-complexes, respectively.

## RESULTS

### Functional characterization of diverse SIV Vpr isolates in human cells

To determine the evolutionary breadth of Vpr-induced DDR signaling, changes to histone marks, and activation of transcription programs, we mined the Los Alamos sequence database for SIV Vpr isolates with diverse phylogenetic lineages. As depicted in **Figure 1A**, we compared 32 SIV Vpr sequences to that of HIV-1_IIIB_ Vpr with a focus on SIV strains previously implicated in cross-species transmission events (Bell and Bedford, 2017). Interestingly, a majority of the SIV Vpr isolates selected for analysis exhibited almost ubiquitous conservation of a C-terminal “SRIG” motif that we previously implicated as being essential for the induction of DDR signaling and epigenetic remodeling (**Figure 1A**, highlighted in purple) (Saladino N, 2026). Isolates highlighted in orange were selected for further analysis because these primate species 1) likely gave rise to HIV-1 Groups M, N and O, 2) likely gave rise to HIV-2, which is also included in our functional analyses, or 3) harbor very little sequence homology to HIV-1_IIIB_ Vpr and thus were predicted to be deficient for inducing DDR signaling (African green money out grouping; AGM) (**Figure 1A**).

Because the genetic sequences of these isolates were incompatible with an HIV-1 provirus, we utilized a previously published construct that harbors a CMV-driven *mCherry-T2A* expression cassette in place of *nef*, which permits independent mCherry and Vpr expression from the same mRNA (**Figure 1B**) (Saladino N, 2026). This allows for an isogeneic infection system wherein all infected cells express HIV-1 Gag, Pol, Tat, and Rev, with Vpr expression being genetically uniform due to the CMV promoter. Virus stocks were generated for each of the indicated Vpr strains and used to determine the functionality of these SIV Vpr isolates in a human expression system.

Due to the fact that there are no commercially available antibodies that recognize this diverse panel of isolates, we utilized two different assays to indirectly assess protein expression and functionality. First, we verified engagement of the host CUL4/DDB1/DCAF1 E3-ubiquitin ligase complex by evaluating relocalization of DCAF1 from the nucleus to the cytoplasm, which is known to occur following Vpr binding (**Figure 1C**) (Le Rouzic et al., 2008; Saladino N, 2026). Comparison of publicly available DCAF1 protein sequences in the NCBI database indicated that DCAF1 for chimpanzee, gorilla, rhesus macaque, green monkey, and sooty mangabey species are identical to that of human. HeLa cells were infected at a low multiplicity of infection (approximately 10% infection rate, which equates to an MOI of ∼0.25) with control or Vpr expressing viruses for 48 hours prior to immunofluorescence microscopy analysis of endogenous DCAF1 localization. As anticipated, all SIV Vpr isolates efficiently induced nucleus-to-cytoplasm redistribution of DCAF1 except for SIV_MNE8_, which exhibited a diminished capacity to relocalize DCAF1 (**Figure 1C**). However, SIV_MNE8_ Vpr was still able to induce significant relocalization compared to control infected cells.

As a second approach to confirm SIV Vpr functionality in human cells, we evaluated the ability of these isolates to deplete well-characterized host factors targeted by HIV-1_IIIB_ Vpr using a live-cell fluorescence-based degradation assay. We generated HEK293T cell lines stably expressing eGFP-tagged HIV-1_IIIB_ Vpr substrates exonuclease 1 (EXO1), uracil DNA glycosylase 2 (UNG2), single-strand selective monofunctional uracil DNA glycosylase 1 (SMUG1), or minichromosome maintenance 10 (MCM10) (**Figure 1D**). Following purifying selection, these cell lines were infected with HIV-1_IIIB_ Vpr or control viruses to evaluate substrate degradation efficiency. Flow cytometric and live-cell fluorescence analyses demonstrated that HIV-1_IIIB_ Vpr induced significant depletion of all four target substrates compared to control infected cells (*i.e.*, loss of eGFP fluorescence in mCherry positive cells). We infected these cell lines with our panel of SIV Vpr isolates and all exhibited degradation activity against at least one substrate except for SIV_RCM312_; however, this isolate was able to induce DCAF1 relocalization as efficiently as HIV-1_IIIB_ Vpr, suggesting that it is functional. Together, these observations indicate that all SIV Vpr isolates function in human cells and are able to engage host cell machinery.

### Evolutionarily distinct SIV Vpr isolates induce DDR signaling, changes to histone marks, and activation of transcription factors

Because Vpr’s proviral function is dominant in myeloid cells, we assessed the ability of these SIV Vpr isolates to induce phosphorylation of the DNA repair marker H2A.X in differentiated THP1 cells (macrophage-like). Phosphorylation of H2A.X (γH2A.X) is a hallmark of DDR activation and is an indicator of the activation status of both ATM and ATR DNA repair kinases. Interestingly, 13 out of 16 SIV Vpr isolates induced significant levels of γH2A.X focus formation in differentiated THP1 cells compared to control infected cells (**Figure 2A**, top). Surprisingly, induction of DDR signaling was not constrained to isolates phylogenetically similar to HIV-1_IIIB_ Vpr, as all four SIV_AGM_ isolates efficiently induced γH2A.X foci (**Figure 2A**, top). To confirm that the lack of DDR activation for SIV_RCM_ and SIV_SMM_ isolates was not due to experiments being carried out in human cells, we also evaluated γH2A.X phosphorylation following infection of African green monkey cells (Vero E6). As depicted in **Figure 2A** (bottom), the trends for γH2A.X phosphorylation in infected Vero E6 cells were similar to those observed following infection of differentiated THP1 cells, suggesting that the lack of DDR activation for SIV_RCM_ and SIV_SMM_ isolates is not due to host species.

**Figure 2.**
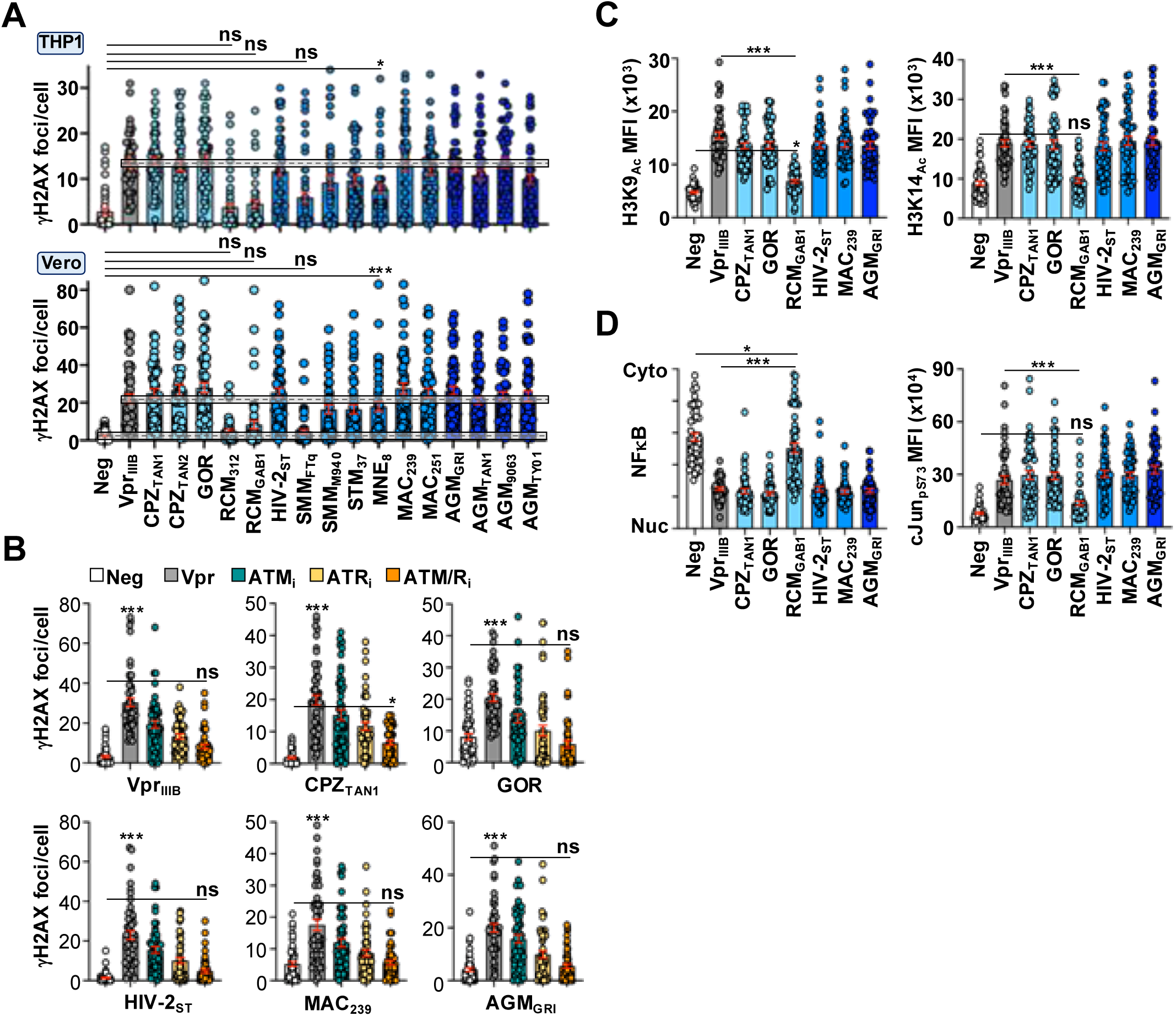
Diverse SIV Vpr isolates induce DDR signaling, epigenetic remodeling, and transcription factor activation. (**A**) Induction of DDR signaling in differentiated THP1 or Vero E6 cells. THP1 cells were simultaneously infected for 48 hours prior to immunofluorescence microscopy to assess γH2AX focus formation. For THP1 differentiation, at the time of infection cells were treated with PMA (100 ng/mL) to induce differentiation for 48 hours prior to immunofluorescence microscopy. For Vero E6 experiments, cells were infected at the time of seeding and prepared for analysis 72 hours post-infection. For quantification, only mCherry positive cells were considered for analysis. Images were obtained using an EVOS M5000 microscope and foci were quantified using Image J software. For foci quantification, we used the “find maxima” imaging processing tool that quantifies foci above a preset threshold. Dashed boxes highlight the mean and standard error mean of γH2AX focus formation in control or HIV-1_IIIB_ Vpr infected cells (n = 50). Analyses performed using a one-way ANOVA; ns, not significant; ***, p < 0.001; *, p < 0.05. (B) Quantification of immunofluorescence microscopy images in differentiated THP1 cells infected with control or Vpr expressing viruses in the presence or absence of vehicle, 10 nM ATM_i_, 10 μM ATR_i_, or combination ATM/ATR inhibitor treatment (n = 50 cells). Cells were infected for 24 hours prior to inhibitor treatment for 24 hours and preparation for immunofluorescence microscopy. Analyses performed using a one-way ANOVA; ns, not significant; ***, p < 0.001; *, p < 0.05. (**C** and **D**) Quantification of immunofluorescence microscopy images of histone marks and transcription factor activation in differentiated THP1 cells infected with control or Vpr expressing viruses (n = 50 cells). Analyses performed using a one-way ANOVA; ns, not significant; *, p < 0.05.

H2A.X phosphorylation is mediated by both ATM and ATR kinases through the activation of DNA double-strand and single-strand break repair pathways, respectively. Since HIV-1_IIIB_ Vpr is known to induce the activation of both repair pathways, we wanted to determine if these SIV Vpr isolates induce H2A.X phosphorylation in a similar manner. To probe this, we treated infected THP1 cells with ATM (AZD1390), ATR (NU6027), or combination ATM/ATR inhibitors 24 hours post-infection and assessed H2A.X phosphorylation at 48-hours post-infection (*i.e.,* a 24-hour inhibitor treatment). All SIV Vpr isolates exhibited similar inhibition profiles compared to THP1 cells infected with HIV-1_IIIB_ Vpr, indicating that both ATM and ATR are contributing to γH2A.X focus formation in SIV Vpr infected cells (**Figure 2B**).

Our group recently established that HIV-1_IIIB_ Vpr-induced DDR signaling leads to epigenetic remodeling and activation of multiple transcription factors associated with HIV-1 promoter activity (Saladino N, 2026). To verify this mechanistic linkage is also conserved amongst diverse SIV Vpr isolates, we assessed the acetylation of two key histone residues associated with transcription activation and the formation of euchromatin as well as the activation of two transcription factors known to influence HIV-1 promoter activity (Bannister et al., 1993; Guo et al., 2011; Lee et al., 2001; McCool and Miyamoto, 2012; Nowak et al., 2008). SIV Vpr isolates capable of inducing γH2A.X foci also induced acetylation of histone 3 amino acid residues lysine 9 (H3K9_Ac_) and lysine 14 (H3K14_Ac_) in differentiated THP1 cells (**Figure 2C**). As a negative control we included SIV RCM_GAB1_, and as anticipated, it was also unable to induce histone acetylation (**Figure 2C**). Next, we investigated the activation of NFκB and cJun transcription factors as they have been previously implicated in DDR signaling and are known to influence HIV-1 promoter activity (Bannister et al., 1993; Kumar et al., 1998; Lee et al., 2001; McCool and Miyamoto, 2012; Nowak et al., 2008; Saladino N, 2026; Varin et al., 2005). As above, all SIV Vpr isolates except RCM_GAB1_ were able to induce nuclear translocation of NFκB and phosphorylation of cJun serine 73 (pS73) in infected THP1 cells. Together, these observations indicate that SIV Vpr-induced DDR signaling also promotes changes to H3K9 and H3K14 acetylation and induces the activation of transcription factors known to be important for HIV-1 promoter activity.

### SIV Vpr isolates utilize convergent and divergent amino acid residues to induce DDR signaling

We and others have demonstrated that Vpr amino acid residues required for inducing DDR signaling are located within the C-terminal domain encompassing an “SRIG” motif (Li et al., 2020; Saladino N, 2026). Interestingly, Vpr isolates derived from RCM_GAB1_ and RCM_312_ both failed to induce DDR signaling yet contain an “SRIG” motif, suggesting an additional protein determinant is required for this activity (**Figures 1A**). To identify additional amino acid residues required for Vpr-induced remodeling of the nuclear environment, we first compared putative Vpr-DCAF protein models for SIV RCM_GAB1_ and AGM_GRI_ Vpr to that of HIV-1_IIIB_ Vpr to identify surface exposed amino acid residues that could be contributing to the activation of DDR signaling. As depicted in **Figure 3A**, the overall protein structures and putative co-complexes of SIV Vpr and DCAF1 closely resembled the HIV-1_IIIB_ Vpr and DCAF1 co-structure (corresponding alpha-fold confidence scores were 0.85/0.85, 0.87/0.88, and 0.67/0.7 for HIV-1, RCM_GAB1_, and AGM_GRI_ Vpr-DCAF co-complexes, respectively) (Wu et al., 2016). Next, we compared HIV-1, CPZ, GOR, and RCM Vpr amino acid sequences to identify residues that were different in both RCM strains (non-functional) compared to the others (functional) and identified 11 polymorphisms of interest (POI) (**Figure 3B**). When all 11 polymorphisms in RCM_GAB1_ were exchanged with the corresponding amino acid resides from HIV-1_IIIB_ (RCM_REV_), we observed a robust induction of H2A.X focus formation that was comparable to HIV-1_IIIB_ control infected THP1 cells (**Figure 3C**). To further define which residues were required for the RCM_REV_ phenotype, we generated additional mutations that partially reverted the HIV-1_IIIB_ residues back to those of wild-type RCM_GAB1_ and assessed DDR activation. Only one reversion mutant exhibited a loss of DDR activation in infected THP1 cells, which encompassed 6 amino acid polymorphisms constrained to the N-terminal half of RCM_REV_ Vpr (**Figure 3C**, RCM_R2_). To confirm these mutations were conferring the DDR activation phenotype, we mutated the corresponding residues in HIV-1_IIIB_ to those of RCM_GAB1_ and as expected, HIV-1_IIIB_ was ablated for DDR activation (**Figure 3D**). Importantly, these Vpr loss-of-function mutants were still capable of depleting SMUG1-eGFP, indicating that these mutants were not compromised for overall function (**Figure 3E**).

**Figure 3.**
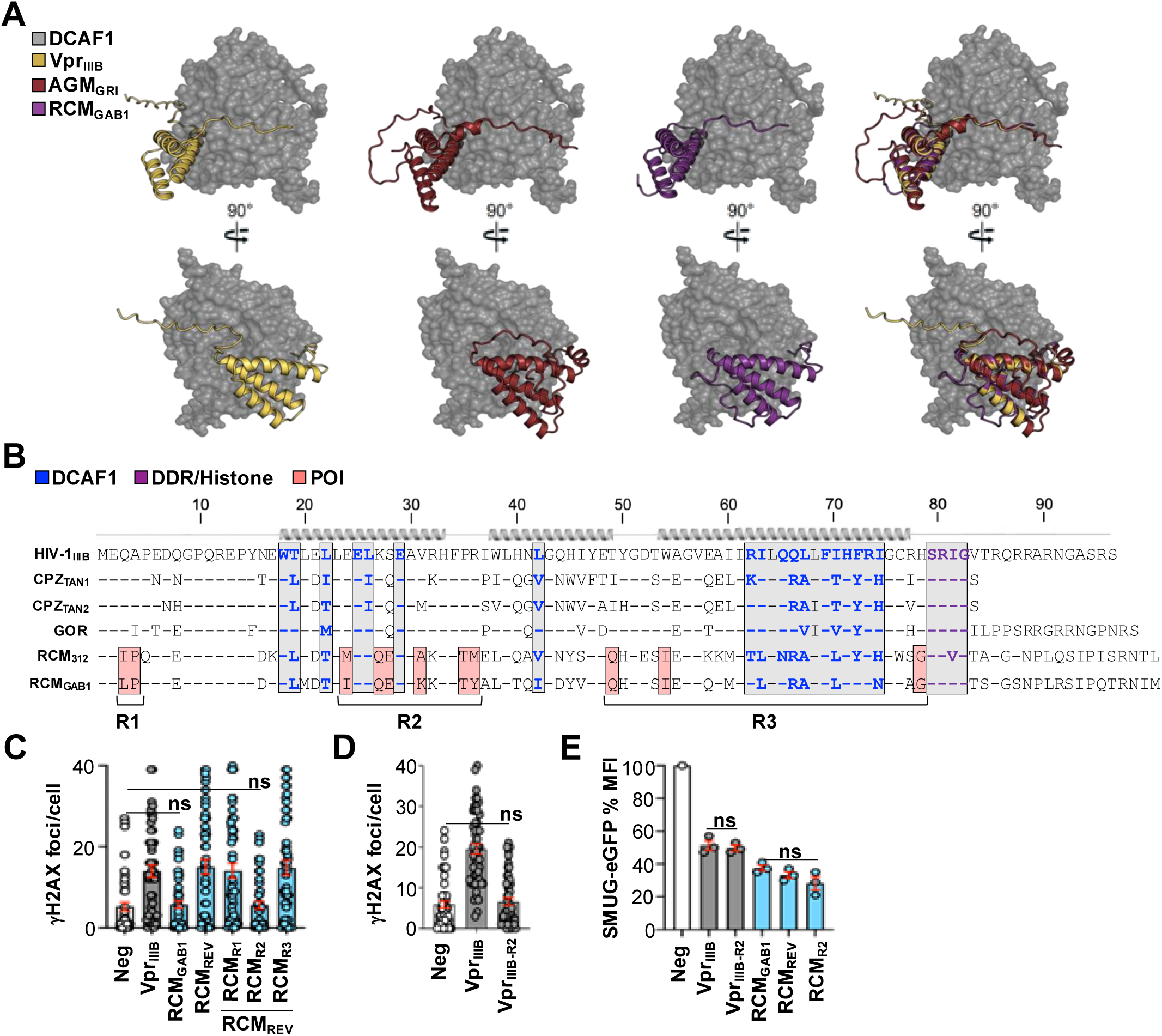
Identification of novel amino acid residues required for HIV-1_IIIB_ Vpr-induced DDR signaling. (A) Predictive modeling of SIV Vpr-DCAF1 co-complexes. SIV Vpr protein modeling was performed using AlphaFold modeling software and overlaid onto a co-crystal structure of HIV-1_IIIB_ Vpr and DCAF1 (PDB: 5JK7). (B) Amino acid alignment of the indicated Vpr isolates. Amino acid residues required for DCAF engagement are colored in blue, previously identified amino acid residues required for induction of DDR signaling, epigenetic remodeling, and transcription activation are colored in purple, and RCM_GAB1_ polymorphisms of interest are colored in salmon. Dashed lines represent amino acid residues that are conserved relative to HIV-1_IIIB_ Vpr. (**C** and **D**) Functional testing of wild-type and mutant HIV-1_IIIB_ and RCM_GAB1_ isolates for their ability to induce H2A.X foci in differentiated THP1 cells (n = 50 cells). R1, R2, and R3 derivatives exchange the amino acid residues highlighted in Figure 3B from wild-type RCM_GAB1_ to those of HIV-1_IIIB_ Vpr. ns, not significant. (**E**) SMUG1-eGFP degradation activity of the indicated Vpr isolates. The SMUG1-eGFP cell line depicted in (Figure 1D) was infected for 48 hours with the indicated viruses prior to being subjected to flow cytometry to quantify the amount of substrate depletion (*i.e.,* quantifying mean eGFP intensity in mCherry positive cells). Mean eGFP fluorescence is depicted to highlight substrate depletion.

Next, we wanted to investigate how the SIV_AGM_ isolates induce DDR signaling even though they share little amino acid similarity to HIV-1_IIIB_ Vpr. Using our putative AGM_GRI_ Vpr-DCAF1 model, we generated single amino acid substitution mutations at surface exposed residues and evaluated H2A.X phosphorylation following infection (**Figures 4A** and **4B**). Interestingly, we identified several mutations that exhibited a diminished capacity to induce γH2A.X foci compared to AGM_GRI_ wild-type infected cells (**Figure 4C**). Importantly, these residues coalesced on the same surface of α-helix 2 (surface outlined in **Figure 4A**), suggesting that they form a functional interface unique from that of HIV-1_IIIB_ Vpr. Moreover, these AGM_GRI_ loss-of-function Vpr mutants exhibited robust degradation activity against SMUG1-eGFP, indicating that they are not globally defective for protein function (**Figure 4D**).

**Figure 4.**
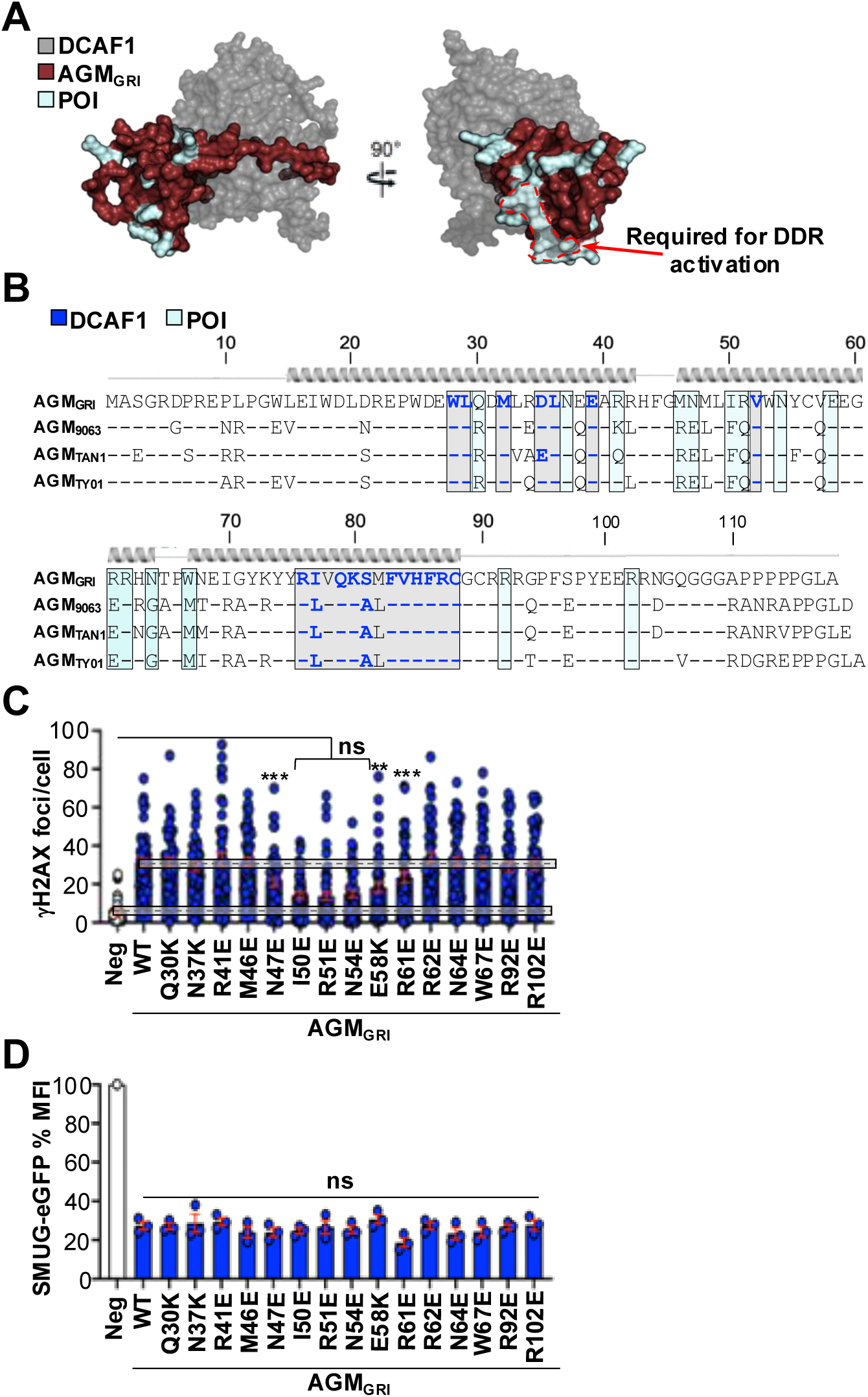
Defining the AGM_GRI_ Vpr surface required for inducing DDR signaling. (**A**) Predictive modeling of a SIV AGM_GRI_ Vpr-DCAF1 co-complex with target surface residues highlighted in cyan. SIV Vpr protein modeling was performed using AlphaFold modeling software and overlaid onto a co-crystal structure of HIV-1_IIIB_ Vpr and DCAF1 (PDB: 5JK7). Residues required for inducing DDR signaling are outlined in a dashed red line. (**B**) Amino acid alignment of the indicated Vpr isolates. Amino acid residues required for DCAF engagement are colored in blue with AMG_GRI_ surface residues of interest highlighted in cyan boxes. Dashed lines represent amino acid residues that are conserved relative to AGM_GRI_ Vpr. (**C**) Functional testing of wild-type and mutant AGM_GRI_ Vpr proteins for their ability to induce H2A.X foci in differentiated THP1 cells (n = 50 cells). ns, not significant; **, p < 0.01; ***, p < 0.001. (**D**) SMUG1-eGFP degradation activity of the indicated Vpr isolates. The SMUG1-eGFP cell line depicted in (Figure 1D) was infected for 48 hours with the indicated viruses prior to being subjected to flow cytometry to quantify the amount of substrate depletion (*i.e.,* quantifying mean eGFP intensity in mCherry positive cells). Mean eGFP fluorescence is depicted to highlight substrate depletion.

## DISCUSSION

While Vpr has been studied extensively, direct cause-and-effect mechanism(s) underlying its proviral function(s) are still incompletely understood. Here, we demonstrate that Vpr’s ability to induce remodeling of the nuclear environment is broadly conserved amongst diverse SIV Vpr isolates. This is noteworthy because 1) recent studies revealed that HIV-1 Vif and Vpu independently antagonize diverse DDR pathways to block the activation of antiviral defenses triggered by abnormal DDR signaling or recognition of viral cDNA (Volcic et al., 2020; Wong et al., 2023), 2) Vpr-induced DDR activation promotes global epigenetic remodeling, activation of diverse transcription programs, and enhanced HIV-1 promoter activity (Saladino N, 2026), and 3), accumulating evidence suggests that this Vpr activity may enhance virus replication through both pre- and post-integration mechanisms, rationalizing why SIV and HIV Vpr isolates would maintain this function over such a broad evolutionary timeframe.

Interestingly, amino acid conservation was not an accurate predictor for DDR modulating activity as several SIV Vpr isolates closely related to HIV-1 Vpr failed to induce a DDR response (*e.g.*, RCM_312_ and RCM_GAB1_). Leveraging these SIV Vpr isolates in combination with mutational and functional analyses revealed a previously uncharacterized set of amino acid residues required for HIV-1 Vpr-induced DDR signaling. Surprisingly, phylogenetically diverse SIV isolates that share virtually no amino acid similarity to that of HIV-1 Vpr were able to induce robust DDR signaling (**Figure 2A**), changes to two histone marks (**Figure 2C**), and activation of transcription factors previously implicated in enhancing HIV-1 promoter activity (**Figure 2D**). Functional conservation of these Vpr activities despite divergent protein sequences suggests that maintenance of DDR modulating activity is potentially advantageous for virus replication. Our phylogenetic analyses indicate that DDR modulation is broadly conserved amongst diverse Vpr isolates, which is somewhat surprising given the limited sequence homology between HIV-1 Vpr and several of the SIV Vpr isolates (most notably SIV_AGM_ strains). However, structural modeling suggests that despite little-to-no amino acid similarity between HIV-1 and SIV_AGM_ Vpr isolates the protein fold is predicted to be remarkably conserved (**Figure 3**). These observations are consistent with previous structural studies that utilized NMR and X-ray crystallography techniques to resolve the structures of Vpr and Vpx isolates from SIV mandrill and rhesus macaque (Schwefel et al., 2015; Wu et al., 2015). Mutagenesis studies indicated that several of the SIV_AGM_ Vpr amino acid residues necessary for DDR induction are polar (Asn_47_ and Asn_54_) or are charged (Arg_51_, Arg_61_, Glu_58_) (**Figure 4**). This is noteworthy given that HIV-1 Vpr likely uses a network of electrostatic interactions to module DDR responses, as N-terminal electropositive amino acid mutations result in hyperactivation of DDR responses, while C-terminal electronegative amino acid substitutions are ablated for DDR activity (Saladino N, 2026). Thus, the combination of structural homology combined with the conservation of surface properties required for DDR activation points toward a mechanism of functional convergence.

It has also been established that some SIV_AGM_ Vpr isolates have the capacity to induce depletion of the myeloid cell-specific restriction factor SAMHD1 (Lim et al., 2012), which is traditionally an activity restrained to the accessory protein Vpx. Vpx is an accessory protein unique to HIV-2 and select SIV lineages that likely arose from a gene duplication event using Vpr as a template (Tristem et al., 1992). Therefore, it is possible that some SIV strains harboring Vpr isolates deficient for modulating DDR activity express a Vpx derivative that can compensate. For example, SIV_RCM_ viruses harbor the *vpx* gene, which could potentially explain why the two Vpr isolates from SIV_RCM_ examined in this study fail to induce DDR responses. Moreover, both Vpr and Vpx package at high levels into nascent viral particles, and intravirion Vpr has been shown to induce remodeling of the nuclear environment prior to provirus integration (Greenwood et al., 2019; Saladino N, 2026). This is noteworthy as our previous study demonstrated that Vpr-induced DDR signaling promotes R-loop accumulation, and recent studies have indicated that HIV-1 preferentially integrates at sites of R-loop formation (Park et al., 2024; Penzo et al., 2025; Saladino N, 2026). Thus, another proviral function of this Vpr activity may be to enhance HIV-1 integration efficiency in terminally differentiated cell populations where the nuclear architecture is more constrained compared to actively proliferating CD4^+^ T cells. Future dedicated studies will be necessary for testing these possibilities.

## DECLARATIONS

### Ethics approval and consent to participate

NA

### Consent for publication

All authors consent to publish this article

### Availability of data and materials

All reagents and protocols used in this study are available upon request to the corresponding author

### Author Contributions

Conceptualization: DJS Funding Acquisition: DJS

Formal Analysis: DJS, ZW, NS, EL, MJ Investigation: ZW, NS, EL, MJ Methodology: ZW, NS, EL, MJ] Project Administration: DJS Writing—original draft: DJS, ZW Writing—review & editing: DJS, ZW, NS, EL, MJ

## Acknowledgement

NA

